# Plasma-derived HIV Nef+ exosomes persist in ACTG384 study participants despite successful virological suppression

**DOI:** 10.1101/708719

**Authors:** Andrea D. Raymond, Michelle J. Lang, Jane Chu, Tamika Campbell-Sims, Mahfuz Khan, Vincent C. Bond, Richard B. Pollard, David M. Asmuth, Michael D. Powell

## Abstract

Human Immunodeficiency Virus (HIV) accessory protein Negative factor (Nef) is detected in the plasma of HIV+ individuals associated with exosomes. The role of Nef+ exosomes (exNef) in HIV pathogenesis is unknown. We perform a retrospective longitudinal analysis to determine correlative clinical associations of exNef plasma levels in ARV-treated HIV+ patients with or without immune recovery. exNef concentration in a subset of AIDS Clinical Trial Group (ACTG) 384 participants with successful virological suppression and with either high (Δ >100 CD4 cell recovery/High Immunological Responders (*High*-IR) or low (Δ ≤100 CD4 cell recovery/ Low Immunologic Responders (*Low*-IR) immunologic recovery was measured and compared for study weeks 48, 96, and 144. CD4 recovery showed a negative correlation with exNef at study week 144 (*r* = −0.3573, \**p*=.0366). Plasma exNef concentration in high IRs negatively correlated with naïve CD4 count and recovery (*r* = −0.3249, \**p* = 0. 0348 (*High*-IR); r =0.2981, \**p*= #0.0513 (*Low*-IR)). However, recovery of CD4 memory cells positively correlated with exNef (r =.4534, *p=.0358) in *Low*-IRs but not in *High*-IRs. Regimen A (Didanosine, Stavudine, Efavirenz) lowered exNef levels in IRs by 2-fold compared to other regimens. Nef+ exosomes persist in ART-treated HIV+ individuals despite undetectable viral loads, negatively correlates with naive and memory CD4 T cell restoration and may be associated with reduced immunological recovery. Taken together, these data suggest that exNef may represent a novel mechanism utilized by HIV to promote immune dysregulation.

## Introduction

The prognosis for patients infected with HIV has improved significantly with the advent of combination antiretroviral therapy (cART) which can lead to suppression of plasma viremia usually associated with marked improvements in CD4+ T cell counts [1, 2]. However, a subset of patients do not experience a robust immune recovery despite viral suppression. While host factors such as age and baseline CD4^+^ T-cells with a naïve phenotype have been observed to be negatively associated with the magnitude of CD4^+^ T-cell count improvement [1, 3–6], virological factors that contribute to these immunologic outcomes are less well defined.

Immune reconstitution can be defined as an increase in the number of peripheral CD4 T-cells to greater than 350-500 cells/dL after 4 years of effective cART [3]. Discordant responses characterized by a lack of immune recovery despite viral suppression occur in 7-39% of participants receiving cART [4–7]. The reasons for this phenomenon are not understood.

The HIV-encoded Negative factor (Nef) has been implicated in pathogenesis based on studies examining HIV-infected long term non-progressors (LTNP) and elite controllers (EC). LTNPs remain asymptomatic for years with stable CD4 counts while ECs control plasma HIV RNA in the absence of ART [12]. Although several hypotheses have been presented to account for the protective factors leading to delayed disease progression, replication defective virions/*nef*-defective virions have been associated with these populations in some cases [9–11]. Recent reports have shown that the viruses within some LTNPs and ECs are actually replication competent and control of HIV may be due to host and/or viral factors that maintain low levels of viremia [11]. HIV-1 Nef may be integral to the control of viral replication and/or T-cell responses in an HIV-infected individual. Early *in vitro* studies have shown that soluble Nef is cytotoxic to T-cells and is released from HIV-infected cells in plasma derived microvesicles that can be detected in HIV-infected patients [13–23]. These studies suggest that Nef does not exist as a soluble protein *in vivo* but instead non-virion associated Nef is found in microvesicles or exosome-like microvesicles (exNef) [7, 8].

Several *in vitro* studies have identified pathogenic activities of soluble Nef including its capacity to reduce surface expression of CD4 and MHC class II, to increase HIV infectivity, to stimulate primary macrophages to release pro-inflammatory cytokines and chemokines, and induce apoptosis in CD4 T-cells [13, 18, 24–27]. We have previously shown that expression of *nef* in HEK293 cells is sufficient to produce exNef, and the resultant exosomes can be absorbed by T-cells and macrophage but only induces apoptosis in T-cells [14]. Taken together, these observations suggest that Nef+ microvesicles/exosome could have an impact on CD4 T-cell recovery. We hypothesize that Nef microvesicles/exosomes are released from HIV infected cells despite successful cART-induced viral suppression and contribute to pathogenesis by impeding CD4-T-cell recovery.

Exosomes range in size from 30–100 nm and are released from hematopoetic (i.e. B-and T-lymphocytes, dendritic cells, monocytes, mast cells, reticulocytes) and non-hematopoetic (i.e. neurons, intestinal epithelial cells, and tumors) cells. Exosomes are detected in physiological fluids such as plasma, urine, malignant effusions, and amniotic fluids. Several functions have been attributed to exosomes/microvesicles including the modulation of cell signaling, cellular homeostasis, intercellular communication, shuttling genetic material, establishment of tissue polarity, regulation of immune responses, and enhancing the site of budding (i.e. HIV and Murine Leukemia Virus) [28–32]. Given the immunomodulatory functions of exosomes, we sought to explore whether exNef may selectively impair CD4+ T-cells recovery during cART.

For this study, Nef concentration was determined in plasma-derived exosomes isolated from a subset of the ACTG-384 cohort with and without immune recovery post-cART. We show that Nef+ exosomes persist and can be detected in study participants with undetectable viral loads even after 144 weeks of therapy. This suggests that exNef production is independent of plasma viral load. Interestingly, we demonstrate that *Low*-IRs have significantly higher levels of exNef compared to *High*-IRs at 144 weeks post-treatment. Recovery of naive CD4 T-cells (CD45RA+, CD62L+) and total CD4 cells negatively correlated with exNef. Overall, we report that Nef+ exosomes are detected in the plasma despite viral suppression and that exNef is negatively associated with changes in naïve and total CD4 count. Taken together these results suggest that exNef may ultimately affect CD4 T-cell recovery and be a biomarker of immune recovery.

## Material and Methods

### Study Population

A subset of cryopreserved plasma samples (n = 240) taken from participants in ACTG 384 were used in this study [1] (Robbins, GK 2003; Shafer 2003). As previously described ACTG 384 was a factorial multi-center randomized controlled trial conducted in the United States and Italy that compared sequential three-drug regimens for treatment of HIV infection. Nine hundred eighty ART-naive HIV-1+ subjects were randomized and treated with stavudine/didanosine or zidovudine/lamivudine with nelfinavir, efavirenz, or both nelfinavir and efavirenz. If virological failure occurred, then participants were placed on another regimen sequentially. In this study, samples from three distinct groups were obtained and assayed. The groups included 1) Treatment failure (TF) defined as those with a virologic failure with a detectable viral load (HIV RNA > 50 RNA copies) what at any time point before 144 weeks, 2). Immunologic responders defined as those with high CD4 improvement of > 100 cells/mm3 (achieved at any point post study initiation) and suppressed viremia (< 50 RNA copies) (High IRs) and 3) Immunologic non-responders defined as those with CD4 improvement of < 100 cells/mm3 and suppressed viremia (Low IRs) (Figure 1). To avoid confounding effects of multiple-treatment regimens, the samples utilized for this sub-cohort were derived from subjects on their first treatment regimen and those with samples from three- or four time intervals (0, 48, 96, and/or 144 weeks).

**Figure 1:**
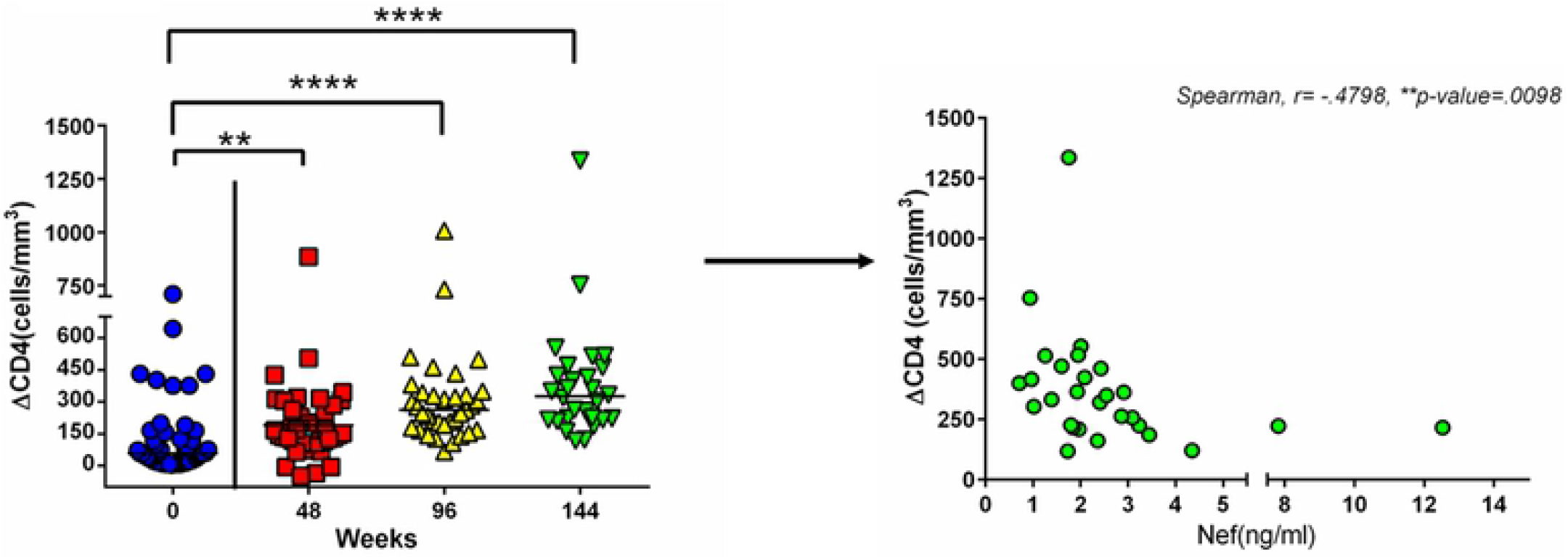
ARV-treated HIV+ patients exhibit differential CD4 T-cell recovery, which negatively correlates with exNef level despite successful viral suppression. **(A)** At study weeks 48 (n=38), 96(n=32), and 144 (n=25) the change in CD4 count (as measured by flow cytometry) was tracked in in the participants with VL >50 RNA copies and compared to baseline CD4 count. High = change in CD4 count >100; Low= change in CD count <100 cells. Statistical significance determined by Kruskal-Wallis test statistic, ***P<.001, Dunn’s Multiple comparison’s **P<.01. **(B)** Plasma-derived exNef concentration negatively correlates with the change in CD4 count. *P<0.05, Spearman r.

### Isolation of plasma-derived exosomes

Plasma exosomes were isolated as previously described [14, 23, 36]. Plasma was pre-cleared by centrifugation at 10,000g for 30 minutes to remove particulates. Microvesicles were pelleted from pre-cleared plasma via ultra-centrifugation for 1 hour at 300,000 x g and then re-suspended in 250 μl of phosphate buffered saline (PBS). Immune complexes within the re-suspended exosomes were removed using acid-dissociation prior to Nef measurement similar to p24 antigen measurements from plasma. Exosomes/high speed pellets (100 μl) were treated with 100 μl of 0.3N hydrochloric (HCl, Sigma) and allowed to incubate for 1 hour at 37 °C. The acid mixture was neutralized with 100 μl of 0.3N sodium hydroxide (NaCl, Sigma) prior to assaying for Nef.

### Nef Enzyme-linked Immunosorbent Assay (ELISA)

Nef concentration in acid-dissociated microvesicle/exosome preparations was measured using a commercially available anti-Nef sandwich ELISA kit (Immunodiagnostics, Bedford, MA) according to the manufacturer’s instructions. Briefly, the neutralized microvesicle preparations were diluted 1:1 with sample diluent (Component C) and added to ELISA plates coated with anti-Nef (Component A). Following 1-hour incubation at room temperature the plates were washed three times with wash buffer (Component B), and 100 μl of anti-Nef-HRP labeled antibody solution (component E) was then added to each well. Plates were incubated for 1 hour at room temperature, washed three times with component B, and developed by adding 100 μl of alkaline phosphatase substrate (component F) per well. Plates were developed for approximately 10 minutes after which 100 μl of stop solution (Component G) was added to each well and the absorbance at 450 nm determined using a Spectramax spectrophotometer.

### Statistical Analysis

Exosomal Nef concentrations were correlated with various immunologic outcomes (immune recovery, change in cell populations, etc.) in each group using the Spearman rank test. The Kruskal-Wallis test was used to compare continuous outcomes between these three groups. The Mann-Whitney rank-sum test or the Dunn’s multiple comparison tests were used appropriately to compare Nef levels in Treatment failures and immunological responders. Statistical analysis was performed using MYSTAT software version 12 (Systat software, Inc. 2007).

## Results

### Baseline characteristics of ACTG384 sub-cohort

This sub-cohort consisted primarily of males (88.4%) (Table 1). Plasma HIV RNA levels at baseline were not significantly different between the three groups (range). Baseline CD4 counts were significantly different between groups (137 cells/mm3 in TF, 60 cells/mm3 in High IRs, 38 cells/mm3 in Low IRs) (as previously described that CD4 count at the initiation of therapy may not play a role in immune recovery [3]. Activated CD4 and CD8 T-cells were not significantly different between the *High*-and Low-IRs groups, either at baseline or post 96 weeks of treatment. (**Table 1**). Other cell subsets B-cells and natural killer (NK) cells were not significantly different between *High*-IRs and *Low*-IRs (data not shown). However, naïve and memory CD4 T-cell subsets were significantly different between *High*-and *Low*_IRs (Table 1) suggesting that these cell subsets are integral to immune recovery.

**Table 1.** Statistical analysis – Intergroup analysis (Column) compared gender and group association using chi-square; ethnicity and group composition at baseline, using Chi-square analysis, #Trend *p-value* <0.1; Median age of groups compared using ANOVA – Kruskal-Wallis, *ns*-not significant; Viral load at baseline between groups at baseline and 96 weeks, Kruskall-Wallis, *ns*-not significant.; Baseline CD4 count, *ns*-not significantly different between groups; Post-96 weeks CD4 count significantly different between Low and High Immunological responders (IR); Kruskall-Wallis, Dunnett’s Multiple comparisons, ***p-value<.001; CD4 count different between baseline and 96 weeks for each group Mann-Whitney ***p-value <.0001; Activated CD4 and CD8 T-cell counts are *ns*-(not significant) between groups at baseline and 96 weeks, Kruskall-Wallis, Dunnett’s Multiple; Naïve CD4 counts is significantly different between high and low IRs at baseline but at 96 weeks differ significantly, Kruskall-Wallis, Dunnett’s Multiple comparisons ***p-value<.001; Intra-group difference in IR memory differ significantly, Mann-Whitney ***p-value <.0001

| Characteristics | Complete subcohort | Treatment Failures | >200 ΔCD4 cell count | <200 ΔCD4 cell count |
| --- | --- | --- | --- | --- |
| sex[n,%] | (n=47) | (n=14) | (n=15) | (n=18) |
| Male | 41,(88.4%) | 10(71%) | 13(87%) | 18(100%) |
| Female | 6,(11.6%) | 4(29%) | 2(13%) | 0 |
| **Age (Baseline)** | 37 | 36 | 37 | 39 |
| **Viral load(log10)** |  |  |  |  |
| Baseline | 5.447 | 4.998 | 5.563 | 5.425 |
| 96 weeks | 1.68 | 3.32 | 1.699 | 1.699 |
| **CD4 Count** | *** | *** | *** | * |
| Baseline | 116 | 252 | 61 | 87 |
| 96 weeks | 330 | 372 | 409 | 253.5 |
| **Activated CD4/CD38/HLA(%)** | *** | *** | *** | *** |
| Baseline | 28 | 21 | 22 | 31 |
| 96 weeks | 7 | 11 | 7 | 6.5 |
| **Activated CD8/CD38/HLA(%)** | *** | * | *** | *** |
| Baseline | 53 | 52 | 57 | 53 |
| 96 weeks | 20 | 29 | 22.5 | 18.5 |
| **Naïve CD4 (cells/mm3)** | *** | ns | *** | *** |
| Baseline | 6.4 | 42.7 | 9.9 | 5.3 |
| 96 weeks | 102.3 | 58.5 | 151.8 | 60.9 |
| **Memory CD4** | *** | ns | *** | *** |
| Baseline | 24.5 | 78.9 | 22.5 | 33.9 |
| 96 weeks | 205.1 | 161.7 | 271.4 | 174.9 |

### CD4 recovery among immunological responders negatively correlates with plasma exNef concentration

ART-induced decrease in viral load (VL) to <50 RNA copies did not necessarily reduce Nef concentration in plasma or result in CD4 recovery for all HIV+ patients within the sub-cohort. We examined study participants receiving anti-retrovirals with successful viral suppression that had CD4 recovery above or below 300 cells/mm^3^ at study week 144 (**Fig 1A**). At study week 144, 40% of the sub-cohort exhibited discordant VL and CD4 cell recovery along with increased exNef level. In the absence of detectable viral replication, exNef could be detected in the plasma and the levels correlated with total CD4 recovery (**Fig 1B**).

### Trend in exosomal Nef correlates with immunological response status

cART initiation reduces viral load to undetectable levels over time but the effects of ART on exNef production is unknown. Cross-sectional reports have demonstrated that Nef can be detected in the plasma in the absence and presence of ART [36]. Since exNef may play a role in HIV immunopathogenesis and impact CD4 recovery we examined the longitudinal changes of exNef in High IRs. Interestingly, exNef was detected in the plasma of these patients despite suppression of viral replication (**Fig 2A**). A retrospective longtitudinal analysis shows that at the initiation of therapy, Nef levels were not significantly different between the *Low*_IRs and *High*_IRs (**Fig 2A**, upper panel). However, by study week 144, the *Low*-IRs had significantly higher levels of plasma exNef than the High IRs (**Fig 2A**, lower panel). The median baseline plasma Nef level in the sub-cohort at 48 weeks was approximately 2.5 ng per 1 ml plasma and by 144 weeks Nef levels increased to almost 5 ng per 1 ml plasma in the Low IRs. In fact *Low*-IRs exhibit an increasing trend in exNef levels from study weeks 48 through 144 while in *High*-IRs exNef level decreased over the study (**Fig 2B**). This finding suggests that plasma exNef may play a role immunological recovery.

**Figure 2:**
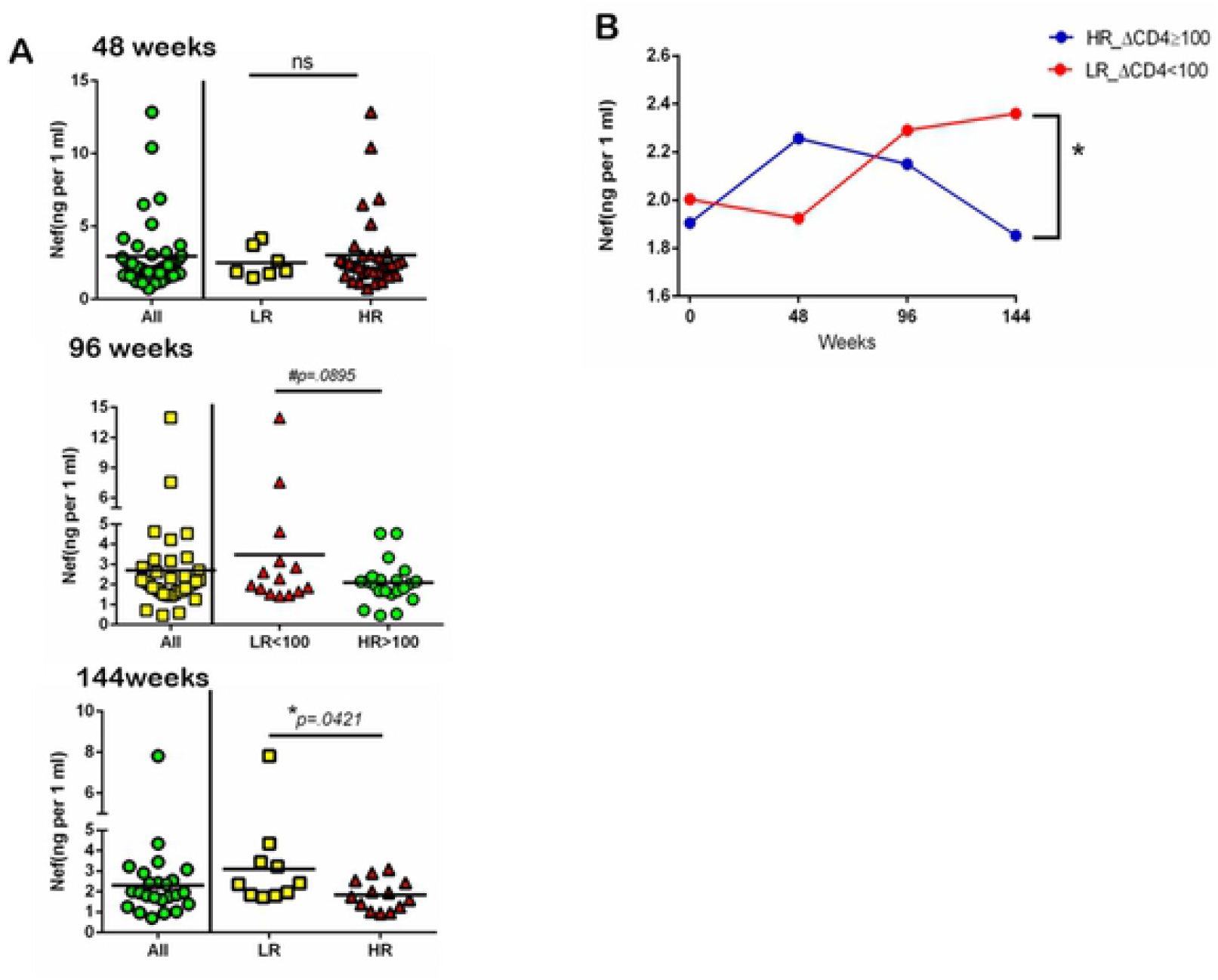
Exosomal Nef is significantly different between low and high immunological responders and trends upward in low immunological responders. **(A)** exNef concentration as measured by anti-Nef ELISA is increased in Low responders at study weeks 48(Low-n=25, High=5, 96(Low-n=16; High-n-14), and 144 weeks (Low-n=10; High n=16). Statistical significance determined via Mann-Whitney, *p-value <0.05, Mann-Whitney. **(B)** High and low immunological responders defined in sub-cohort defined by change in CD4 count from baseline. High – change ≥100 (cells/mm^3^); Low change <100 cells/mm^3^) (B) Median Nef concentration in exosomes isolated from plasma of subjects with Treatment Failure (n=10) or Success (with high, and low CD4 recovery; n=30) subjects. Nef quantified by anti-Nef ELISA.

### Naïve CD4 count and recovery inversely correlate with exNef

We then investigated whether Nef+ exosome levels correlated with recovery of naïve CD4 cell counts specifically. Immunological recovery (as defined in Methods) 48 weeks post treatment and close to 90% (18 out of 20) by week 96 of treatment (**Fig 3A**). However, none of the Low-IRs exhibited increases in CD4 T-cell count close to 350 cells/mm^3^ by 144 weeks (**Fig 3A**). Although naïve T-cells do recover in both *High-* or Low-IR the IRs have appreciably less naïve CD4 cells than the IRs 96-and 144 weeks post treatment initiation (**Fig 3C**). Notably the changes in both CD4 T-cell count and CD4 naïve T-cells negatively correlated with the Nef concentration in plasma-derived microvesicles (**Fig 3, B and D**) suggesting that *in vivo* microvesicular Nef may be associated with immune recovery/ CD4 T-cell rebound.

**Figure #3:**
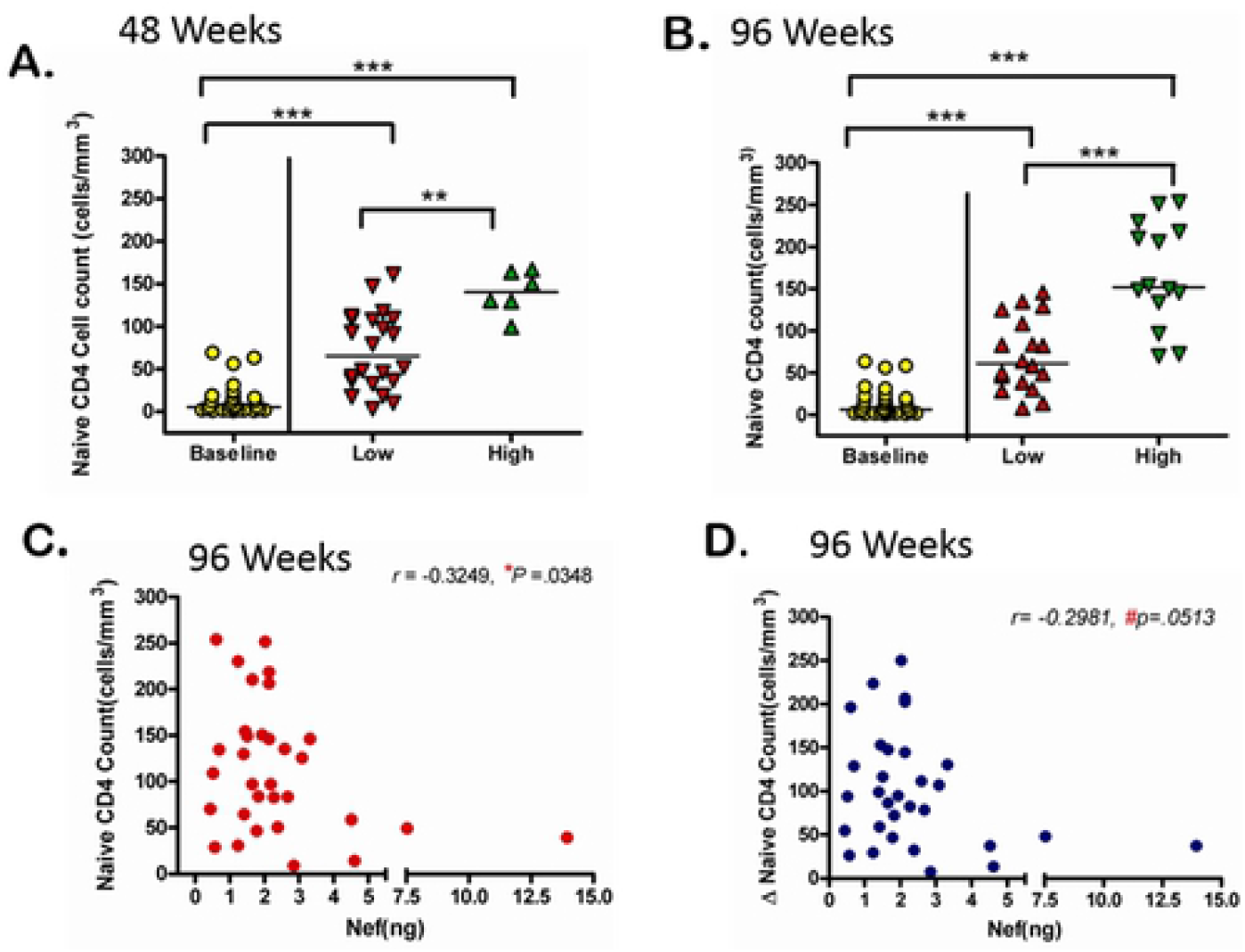
Naïve CD4 counts inversely correlate with exNef. **(A)** Naïve counts (as measured by flow cytometry) significantly higher in the High_IR group at study week 48 and **(B)** study week 96. (C) Nef-level inversely correlates with both Naïve CD T-cell count and the (D) change in Naïve CD4 count post ARV-treatment. Correlation determined by Spearman r, *p-value<.05, #p-value <.1 (trend). Statistical significance determined by One-Way ANOVA Kruskal-Wallis test statistic and Dunn’s Multiple comparison * P-value<0.05, **p-value<.01, and ***p-value<.001.

### Memory CD4 T-cells are significantly different in high low and responders

Given the role of memory cells in immunological recovery, we sought to determine whether exNef also impacted CD4 memory cell recovery. Interestingly at weeks 48 and 96 post-therapy *High*-IRs exhibited appreciably more CD4 memory cells than the *Low*_IRs (**Fig 4**, panel A). The degree of CD4 increase directly correlated with exNef in the Low_IRs (**Fig 4**, panel B), suggesting that CD4 memory cells could be one of the sources of exNef during anti-viral suppression.

**Figure #4:**
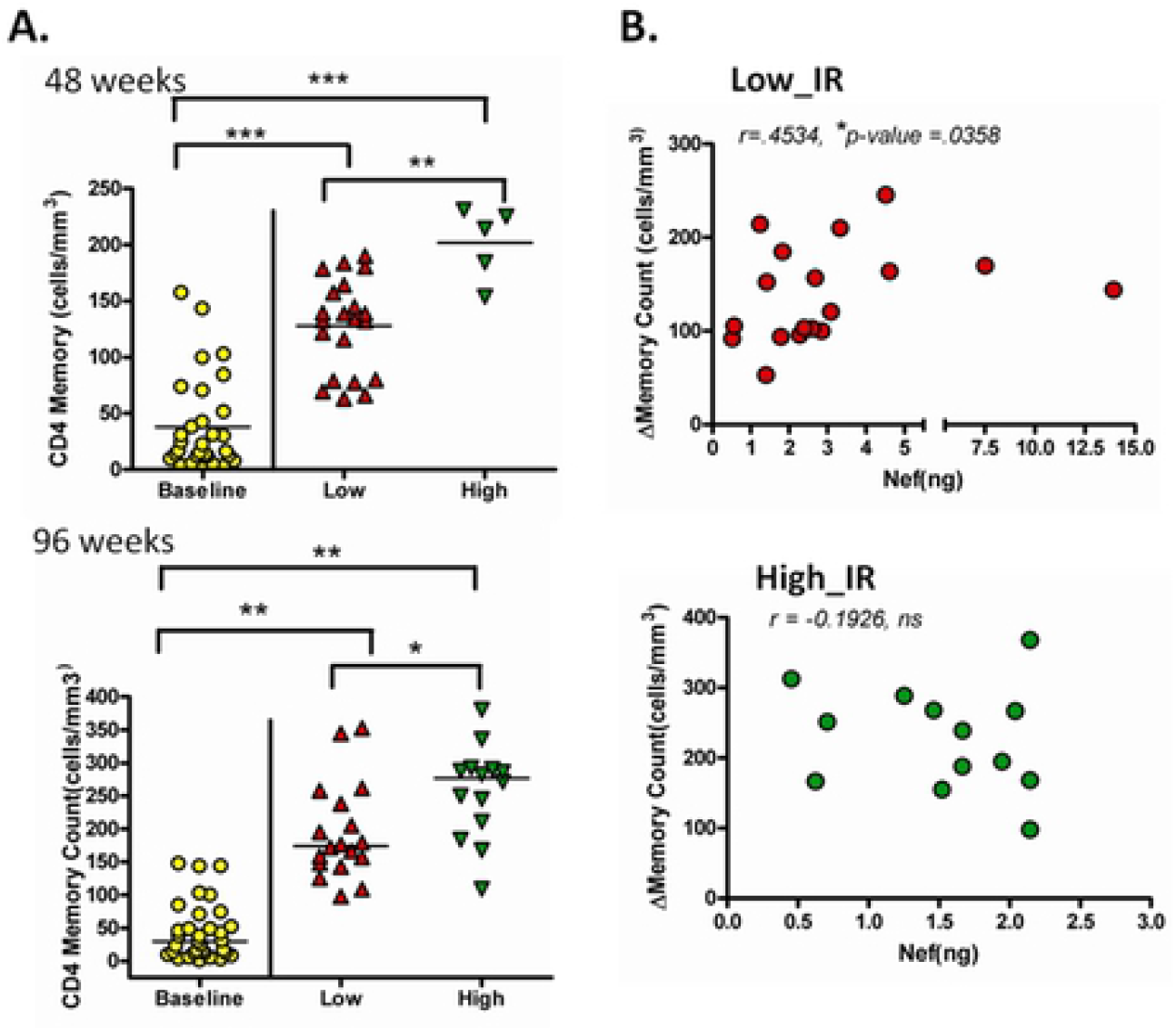
Memory CD4 T-cells are significantly different in high low and responders. **(A)** CD4 memory cell counts (as determined by flow cytometry) in high-immunological responders are significantly increased compared to low responders at study weeks 48 (Low-n=25; High-n=5); and 96 (High-n=14; Low-n=16). Significance from baseline determined via Kruskal-Wallis, intergroup differences compared via Dunn’s Multiple comparison *p-value<0.05, **p-value<.01, and ***p-value<.001. **(B)** The change in CD4 memory count from baseline(n=33) to study week 96 (n=30) directly correlates with exNef level in low-immunological responders (n=18). Correlation determined using Spearman r, *p-value<0.05, ns=not significant.

### PI-sparing regimen significantly reduced plasma exNef

ACTG 384 was a prospective double-blinded study using a factorial design to compare sequential three-drug regimens. Study arms are depicted in **Table 2**. Basically, two NRTIs zidovudine and lamuvidine or didanosine and stauvidine followed by either efavirenz or nelfinavir were compared (**Fig 5A**, upper panel). Previous results from the ACTG-384 cohort demonstrated that the combination of zidovudine, lamivudine, and efavirenz lead to the shortest time to viral suppression suggesting that this combination was the most efficacious combination [2]. However, we are still able to detect exNef in the sub-cohort of ACTG384 participants with successful viral suppression. If exNef negatively affects immunological recovery, then we must identify a treatment regimen that suppresses both viral replication and exNef release. To determine how treatment regimen impacts exNef production we stratified participants by treatment regimen and compared their respective exNef level. By weeks 96 and 144 exNef was reduced 2-fold only in participants (with undetectable viral load) that received regimen A (Didanosine, Stauvidine and Efavirenz) (**Fig 5**, lower panel). This suggests that drug regimen may also dictate exNef levels and that PI-sparing regimens reduce both viral load and exNef level.

**Figure #5:**
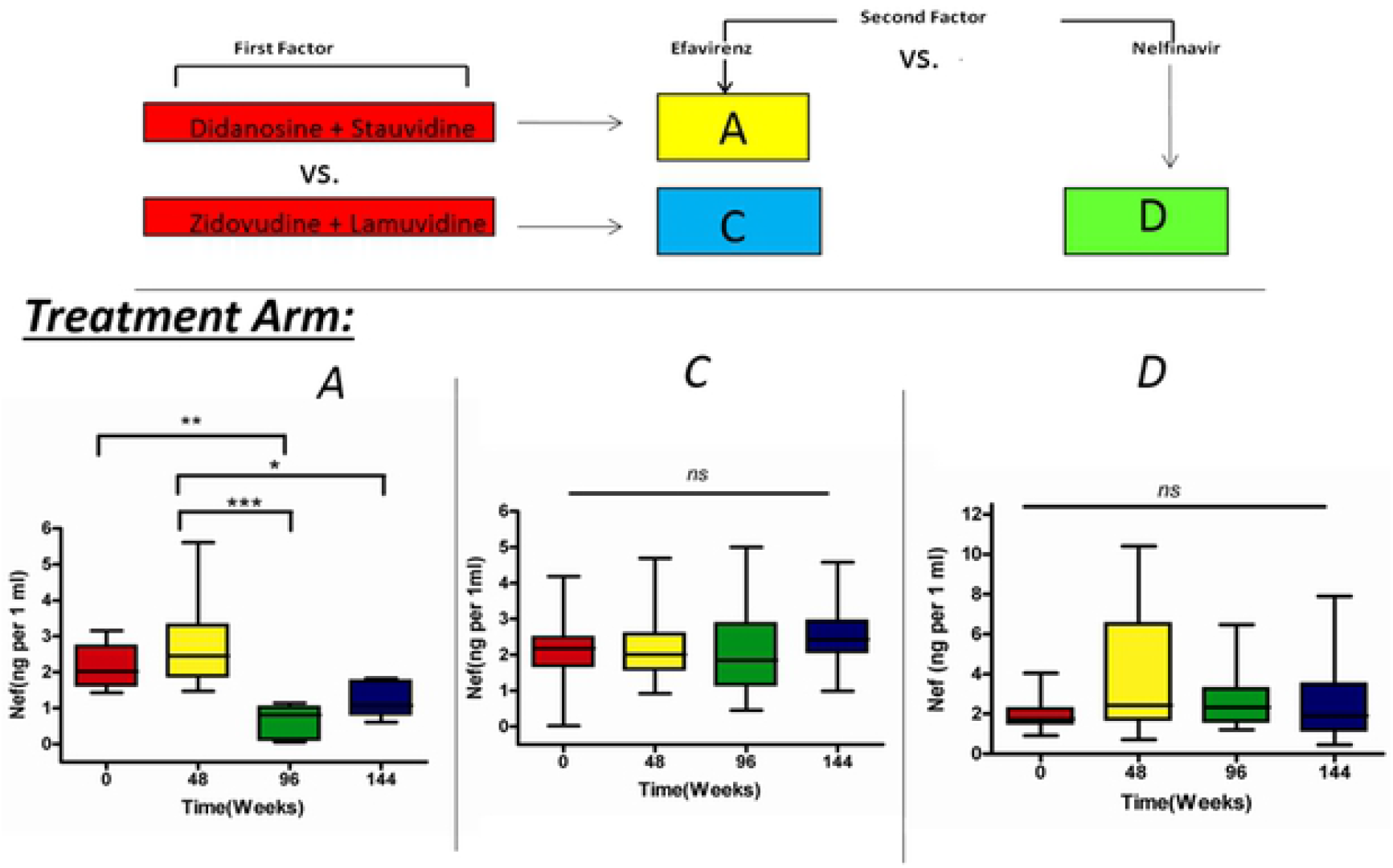
PI-sparing regimen significantly reduced plasma exNef by study weeks 96 and 144. (Upper panel) Schematic of drug regimens administered to sub-cohort in the first tier of the ACTG384 study. Participants on regimens A (n=5) or C (n=10) received efavirenz with DDI/4TC or ZDV/4TC while those given regimen D(n=7) received ZDV/4TC and Nelfinavir(**Lower Panel**). Longtitudinal comparison of plasma exNef level as measured by anti-Nef ELISA in participants taking regimen A,C, and D. Statistical significance determined via Kruskal-Wallis, and Dunn’s multiple comparison *p-value <0.05, **p-value<.01, ***p-value<.001, and ns=not significant.

**Table 2:** Study arms of ACTG384. Drug regimen groupings as described by Smeaton et al 2001 followed by the number of participants (in parentheses) within regimen group.

| Initial | Initial NNRTI | Initial PI | Initial PI +NNRTI |
| --- | --- | --- | --- |
| NRTI Factor | Efavirenz<br>(EFV) | Nelfinavir<br>(NFV) | NFV +<br>EFV |
| DDI+D4T | A (5) | B (3) | E (11) |
| ZDV /3TC | C (10) | D (7) | F(14) |

## Discussion

This study shows that exNef is detected in the plasma of cART-treated HIV+ patients with successful viral suppression. Immunological recovery does not occur in 40% of cART-treated HIV+ patients. Understanding the factors involved in poor immunological recovery could lead to the development of novel therapeutics that inhibit viral replication while promoting immunological recovery. We posit that exNef may represent a novel mechanism utilized by HIV to promote viral replication in resting CD4 T-cells. The impact of exNef on HIV pathogenesis and CD4 T-cell recovery during cART is unknown. Additionally, we demonstrate that exNef level was not only elevated in cART-treated HIV+ patients with low immunological recovery but also negatively correlated with CD4 count recovery in these participants. This clearly suggests a role for exNef in immunological recovery.

Plasma exNef could theoretically determine immune recovery potential. Detection of Nef microvesicles is not unexpected since several studies have shown that in order to lower surface expression of CD4 and MHC class II Nef interact with component of the endocytic and exocytic machinery [13, 22]. Nef has no enzymatic activity and functions primarily as an adaptor within an infected cell. The function of extracellular Nef is unclear. However, several *in vitro* studies have demonstrated that extracellular soluble Nef is cytotoxic to cells, induces cytokine and chemokine release from macrophage activation, increases viral infectivity and alters innate immune signaling pathways [13, 18, 27, 37–40]. We know from *in vitro* and *ex vivo* studies that Nef is released in exosome-like vesicles from nef-transfected and HIV-infected cells and that these vesicles are detected in the plasma of HIV+ patients [14, 23]. Although the biological role of Nef+ exosomes is still unknown we expected exNef to have similar cellular and functional affects ascribed to soluble Nef.

It has recently been reported that released Nef microvesicles/exosomes similar to soluble Nef can trigger activation induced cell death (apoptosis) in peripheral blood leukocytes thus promoting the depletion of CD4+ T-cells [21]. Another report indicated that Nef induces massive secretion of microvesicle clusters in HIV-infected T-cells and that extracellular Nef is then passed to uninfected bystander by cell-to-cell contact via an ERK-1/2 dependent mechanism [41]. Other studies have shown that extracellular Nef vesicles are taken-up/absorbed by T-cells and that these released Nef microvesicles cause activation-induced cell death in primary peripheral blood leukocytes [14, 21]. Primary leukocytes exposed to exNef as observed with soluble Nef release chemokines MIP-1a and MIP-1b (unpublished result).

Here we show that plasma-derived microvesicles/exosomes from participants within the ACTG384 cohort contain Nef even in the presence of successful HAART therapy and are elevated in low immunological responders. This suggests that exNef may play a role in preventing sufficient CD4 T-cell recovery thereby promoting immunodeficiency despite successful HAART outcomes. Taken together, these studies show that extracellular Nef exosomes function similar to soluble Nef in that exNef is absorbed by T-cells, induce apoptosis and may play a role in discordant successful viral suppression and CD4 T-cell recovery.

Most interestingly Nef+ microvesicles/exosomes persist and are detectable over a 144-week period suggesting that exNef may simply be a product of chronic HIV infection. We observed that significant reduction of viral load correlated with an increase in exNef level suggesting that exNef may regulate viral replication and/or cell function. These findings are novel in HIV-pathobiology but may not be a unique to HIV. Indeed, recent reports demonstrated that EBV-infected nasopharyngeal cells release inhibitory exosomes containing the EBV-encoded protein latent membrane protein-1 (LMP-1) [42]. These LMP-1+ exosomes inhibited T-cell activation and anti-EBV immune responses [42]. So exNef released from HIV-infected cells may have a similar effect on anti-HIV immune responses.

Aside from the ACTG384, several studies have shown HIV+ patients with successful virological responses to HAART yet incomplete CD4 recovery have increased mortality [43–45]. Age and prolonged periods of immunodeficiency prior to successful HAART are risk factors for incomplete/insufficient CD4 T-cell recovery [46]. We provide evidence in this study that plasma derived Nef+ microvesicles may be associated with immune recovery. We detected increased in exNef only in *Low*_IRs. In these *Low*-IRs exNef concentration correlated negatively and positively with the recovery of CD4 Naïve and memory T-cell counts, respectively. This result suggests that naïve CD4 cells are a potential exNef targets while the CD4 memory cells could be a source of the Nef+ exosomes.

Treatment regimens not only directly impact viral replication but also appear to affect the generation of Nef+ exosomes. PI-sparing HAART regimens (e.g. Regimen A: Didanosine, Stauvidine, and Efavirenz) reduced both viral load and exNef level. This result suggests that treatment regimen may dictate exNef level and exNef in turn may be developed as a prognostic indicator of CD4 immune recovery during HAART.

We posit that the increased level of Nef+ exosomes early in *High*_IRs (by week 48) are indicative of reduced virus production and suggests that in the absence of productive viral replication HIV-infected cells release more Nef+ exosomes. If these exosomes affect T-cell activation or viability, then this may impact immune recovery. By 144 weeks however, exNef is significantly reduced in *High*-IRs participants relative to *Low*-IRs participants. Taken together our study suggests that extended use of combination therapy lacking a HIV protease inhibitor impairs the release of Nef+ exosomes while NRTI/NNRTIs have no long-term effects on exNef.

Overall, these data also suggest that increased Nef levels maybe a double-edged sword – in terms of viral suppression-high exNef is associated with decreased viral load but in regard to immune recovery-elevated exNef is associated with reduced CD4 T-cell rebound/immune recovery. Recently, regions within Nef important for secretion have been identified. Disruption of the secretion modification region (SMR) within the Nef gene appears to abolish its release in microvesicles/exosomes [61]. Peptide-targeted disruption of the SMR also inhibited Nef release suggesting that SMR peptides could be developed as a therapeutic agent.

Ultimately, we provide evidence of exNef as a novel virological factor contributing to the dissociation of viral load and immunological recovery. Since, drug regimens can alter exNef levels, therapies need to be developed that both successfully lower viral load and Nef+ exosomes levels. Our findings also suggest that clinicians could monitor exNef level along with CD4 T-cell count in patients undergoing ARV-treatment to assess the effectiveness of therapeutic regimens.

## Acknowledgements

We would like to acknowledge the assistance of Dr. Gale Newman (Morehouse School of Medicine). We also acknowledge the support of RCMI core facilities at Morehouse School of Medicine (U54MD007602), the Emory Center for AIDS Research (P30AI050409), and the NIH AIDS Reagent program. We are grateful for funding support from grants S06-GM08428, R21 A1060370, and R21 NS105577). Research reported in this publication was supported by the National Institute of Allergy and Infectious Diseases of the National Institutes of Health under Award Number UM1AI068634, UM1 AI068636 and UM1 AI106701. The content is solely the responsibility of the authors and does not necessarily represent the official views of the National Institutes of Health”.

